# The pre-mRNA Splicing Modulator Pladienolide B Inhibits *Cryptococcus neoformans* Germination and Growth

**DOI:** 10.1101/2025.03.06.641977

**Authors:** Sierra L. Love, Megan C. McKeon, Henrik Vollmer, Joshua C. Paulson, Nanami Oshimura, Olivia Valentine, Sébastien C. Ortiz, Christina M. Hull, Aaron A. Hoskins

## Abstract

*Cryptococcus neoformans* is an opportunistic fungal pathogen responsible for life-threatening infections, particularly in immunocompromised individuals. The limitations of current antifungal therapies due to toxicity and the emergence of resistance highlight the need for novel treatment strategies and targets. *C. neoformans* has an intron-rich genome, and pre-mRNA splicing is required for expression of the vast majority of its genes. In this study, we investigated the efficacy of a human splicing inhibitor, pladienolide B (PladB), as an antifungal against *C. neoformans.* PladB inhibited growth of *C. neoformans* in liquid culture and spore germination. The potency of PladB could be increased by simultaneous treatment with either FK506 or clorgyline. This combination treatment resulted in significant reductions in fungal growth and prevented spore germination. Transcriptomic analysis revealed that PladB inhibits splicing in *C. neoformans* and results in wide-spread intron retention. In combination with FK506, this resulted in down-regulation of or intron-retention in transcripts from processes vital for cellular growth including translation, transcription, and RNA processing. Together, these results suggest that targeting RNA splicing pathways could be a promising antifungal strategy and that the effectiveness of splicing inhibitors as antifungals can be increased by co-administering drugs such as FK506.

**IMPORTANCE:** Fungal infections, like those caused by *Cryptococcus neoformans*, can turn deadly for many patients. New treatments and therapeutic targets are needed to combat these pathogens. One potential target is the pre-mRNA processing pathway which is required for expression of nearly all protein-coding genes in *C. neoformans.* We have determined that a pre-mRNA splicing inhibitor can inhibit both *C. neoformans* growth and germination and that the potency of this drug can be increased when used in combination with other molecules. This work provides evidence that targeting steps in pre-mRNA processing may be an effective antifungal strategy and avenue for development of new medicines.

## INTRODUCTION

Fungal infections are a serious global health threat, especially as the number of immunocompromised individuals rises (Fisher et al., 2022). Worldwide, invasive fungal infections are estimated to cause over one million deaths annually (Bongomin et al., 2017). A major contributor to this mortality rate, especially among people with HIV/AIDS, is the pathogenic yeast *Cryptococcus neoformans* (Rajasingham et al., 2022). Human disease occurs when *C. neoformans* is inhaled from the environment and then disseminates from the lungs to cause fatal meningoencephalitis. Treatments for cryptococcosis and other invasive fungal diseases have included conventional antifungal therapies, such as polyenes, azoles, allylamines, and echinocandins (Lee et al., 2023). However, these treatments often come with significant toxicity and limited efficacy. Compounding these issues, many fungal pathogens have developed drug-resistance mechanisms, resulting in overall mortality rates from invasive fungal disease averaging ∼50% in the United States and higher in other parts of the world (Denning, 2024). These treatment limitations and the consequential toll on patients underscore the urgent need for novel antifungal drug development strategies.

A potential avenue for targeting fungal pathogens is by modulating pre-mRNA splicing, a process essential for normal cellular function. In *C. neoformans*, genes, on average, have 5.7 introns, and nearly all expressed protein-coding genes must be spliced (Gonzalez-Hilarion et al., 2016; Grützmann et al., 2014; Janbon, 2018). Several splicing modulators have been discovered for the human splicing machinery (Effenberger et al., 2017). In human cells, these compounds can be very potent and change splicing outcomes at nM concentrations. Often, these are described as splicing modulators, rather than inhibitors, because they can show highly idiosyncratic effects on splicing. For example, some splicing events can be completely unimpacted by a drug, whereas others in different genes can be completely inhibited under the same conditions (Vigevani et al., 2017).

To date, many of the most effective splicing modulators bind the same pocket of the essential splicing factor SF3B1, which is highly conserved across eukaryotes. These drugs block splicing by competing with the intron branch site RNA for binding to SF3B1 and the U2 small nuclear ribonucleoprotein (snRNP) (Cretu et al., 2018). The drug binding pocket of SF3B1 may be an attractive target for antifungal drug development. Despite its conservation, some SF3B1 orthologs, like that in *Saccharomyces cerevisiae*, are insensitive to human splicing modulators (Carrocci et al., 2018). This suggests that it is possible to inhibit SF3B1 and pre-mRNA splicing with drugs in a species-specific manner. Targeting SF3B1 in fungal pathogens like *C. neoformans* could disrupt gene expression and represent a novel approach for antifungal therapy.

In this study, we investigated the antifungal potential of a splicing modulator (pladienolide B, PladB) in *C. neoformans*. We found that PladB can inhibit both yeast growth and spore germination, and its potency can be significantly increased by addition of either FK506 or clorgyline. Transcriptomic analysis confirmed that PladB causes intron retention and splicing modulation in *C. neoformans* but of only a subset of introns. This analysis also revealed that PladB, in combination with FK506, downregulated or changed the splicing of genes essential for transcription, RNA processing, and translation, providing insights into the mechanisms of growth inhibition. Combined, our results show that a splicing modulator can have potent antifungal activity and suggest novel therapeutic strategies for combating human fungal pathogens.

## MATERIALS AND METHODS

### Strains

*Cryptococcus neoformans* strains (KN99α, JEC20, and JEC21) were handled with standard techniques as described previously (Frerichs et al., 2021; Kwon-Chung et al., 1992).

### Growth Assays

KN99α yeast were struck out on yeast peptone dextrose (YPD) agar plates and incubated at 30℃ for three days. YPD liquid cultures were then inoculated and incubated at 30℃ overnight. Yeast were diluted to an OD_600_ = 0.05 (OD 0.05) and grown for 4.5h at 30℃ with shaking. Yeast were then diluted to 0.10 OD, and 100 µL were dispensed into 96-well clear round-bottom plates (Costar, Corning inc, Kennebunk, ME, ref no. 3788) that contained the compound of interest or dimethyl sulfoxide (DMSO; 1% v/v final) carrier. The plates were then incubated at 30℃ with shaking, and the OD_600_ was measured at regular intervals over the course of 24 h.

### Stamping Assays

KN99α was struck out on YPD plates and incubated at 30℃ for three days. YPD cultures (25 mL) were then inoculated and grown overnight at 30°C with shaking. Cells were diluted to OD 0.05, incubated for 4.5h, diluted again to OD 0.1, and then dispensed into 96-well clear round-bottom plates containing the compounds(s) of interest. Plates were then incubated at 30℃ with shaking for 24 h. Yeast were then isolated, centrifuged (5000 rpm, 1 min), and resuspended in 100 µL of 10% (v/v) glycerol. Diluted cultures were then stamped onto YPD plates and incubated at 30℃ for three days before imaging.

### Spore Isolation

Spores were isolated from *Cryptococcus neoformans* serotype D (*C. deneoformans)* crosses as described previously (Frerichs et al., 2021). In brief, JEC20 and JEC21 yeast were incubated at 30°C on YPD plates for 2 days, then mixed in a 1:1 ratio in phosphate buffered saline (PBS), and plated in 10 µL spots on V8 juice agar (5% w/v agar, 5% v/v V8, 0.5 g/L potassium dihydrogen phosphate pH 7.0) plates. Crosses were maintained at 22°C in the dark for 5 days. Crosses were resuspended in a 75% (v/v) Percoll solution (in PBS) and centrifuged (21,410 x *g*, 25 min, 4°C). Spores were isolated using a needle puncture (21-gauge, Fisher Scientific, Hampton, NH, UGA, ref no. 1484092), washed with PBS and stored at 4°C until use. Spores were counted using a using a hemacytometer.

### Quantitative germination assay

Spore germination was analyzed using a quantitative germination assay as described previously (Barkal et al., 2016). Briefly, populations of pure spores were plated into 384 well plates (Thermo Scientific 142762) at ∼10^5^ cells per well. At t = 0, a nutrient source (Synthetic Defined (SD) medium plus 111 mM glucose) ± inhibitor was supplied to initiate germination. Pladienolide B was included at 30 µM, FK506 at 50 µM, and clorgyline at 50 µM. Cells were imaged every 2 h for 16 h. Cell size and shape were quantified using a custom script in ImageJ and MatLab.

### Combination synergy screens

KN99α yeast were grown and diluted into 96-well plates as described above, containing combinations of the drugs of interest or 1% (v/v) DMSO. Plates were incubated at 30℃ with shaking for 24 h after which OD_600_ values were recorded. Replicates were averaged and compared to the average of the DMSO control samples to calculate the percent growth inhibition. SynergyFinder+ (Zheng et al., 2022) was used to produce combination plots and synergy heatmaps, using scoring criteria in which combinations with Bliss synergy scores below -10 are antagonistic, scores between -10 and 10 are additive, and scores above 10 are synergistic (Malyutina et al., 2019; Zheng et al., 2022).

### RNA Isolation

KN99α was struck out on a YPD plate and incubated at 30°C for three days. Colonies (4 biological replicates) were then used to start separate liquid cultures (10 mL in YPD) and allowed to grow overnight at 30°C. Cells were diluted to OD 0.05 in 10 mL of YPD and grown for 8h at 30°C. Cells were then diluted to OD 0.5 in 1mL of YPD and incubated with 10 mM FK506, 10 mM PladB, both 10 mM FK506 and 10 mM PladB, or 1% (v/v) DMSO for 2 h at 30°C. Cells were isolated by centrifugation at 4000 rpm for 7 min, and pellets were flash-frozen in liquid nitrogen. Total RNA was extracted from *C. neoformans* cell pellets as described previously (Collart & Oliviero, 1993). Isolated RNA was then DNAse treated and further purified using a Monarch® RNA Cleanup kit (New England Biolabs).

### RNA Library Prep and RNA-sequencing

RNA quality control, library preparation, and sequencing were conducted by GENEWIZ (Azenta Life Sciences). Library prep involved polyA selection, cDNA synthesis, and adapter ligation. Sequencing was completed on an Illumina® NovaSeq^TM^ platform with a read depth of ∼40-50 million paired-end reads per sample. Data are available via the NCBI Sequence Read Archive (BioProject ID PRJNA1226125).

### RNA-seq Data Analysis

Quality control of RNA-seq reads was performed using FastQC. Adapter trimming was carried out with Cutadapt 4.9 (Martin, 2011). Trimmed reads were then aligned to the *C. neoformans* var. grubii H99 genome sequence (NCBI RefSeq ID: GCF_000149245.1) using STAR 2.7.7a (Dobin et al., 2013). A minimum of 40 million reads per strain per replicate was obtained. Gene-level read quantification was performed using the R package Rsubread v2.20.0 (Liao et al., 2019). Differential gene expression analysis was conducted with DESeq2 v1.46.0 (Love et al., 2014), and differentially expressed genes (DEGs) were identified using thresholds of log_2_ fold change > 1 or < -1 and an adjusted *p*-value < 0.05. Alternative splicing (AS) analysis was performed using SpliceWiz v1.8.0 (Wong et al., 2023). Differentially spliced events were identified based on a change in percent spliced-in (ΔPSI) > 0.1, a false discovery rate (FDR) < 0.05 and a log_2_ fold change > 1 or < -1. Functional enrichment analysis of significant DEGs and alternatively spliced genes was conducted using FungiDB, with enriched Gene Ontology (GO) terms identified against the *C. neoformans* var. grubii H99 genome background.

## RESULTS

### Pre-mRNA Splicing Modulators Inhibit *C. neoformans* Growth and Germination

To determine if *C. neoformans* might be susceptible to splicing modulation by SF3B1-targeting compounds, we analyzed conservation of the SF3B1 drug-binding pocket relative to drug-sensitive (*H. sapiens*) and drug-resistant (*S. cerevisiae*) orthologs (**Fig. 1A**). A previous study demonstrated that a N747V substitution in *S. cerevisiae* SF3B1 (relative to human SF3B1) results in drug sensitivity that is significantly enhanced if the protein also contains a L777N substitution (Hansen et al., 2019). This suggests that SF3B1 proteins harboring hydrophobic and hydrophilic amino acids at positions corresponding to *S. cerevisiae* 747 and 777, respectively, are sensitive to drugs that bind this pocket. Proteins with the opposite features are likely resistant. *C. neoformans* contains a hydrophobic amino acid at the orthologous position of amino acid 747 (V924 in *C. neoformans* SF3B1) and a hydrophilic amino acid at position 777 (S954). Correspondingly, this suggests that *C. neoformans* should be sensitive to SF3B1-binding splicing modulators. These features are not shared among all pathogenic fungal organisms since the *Candida albicans* SF3B1 amino acid sequence more closely resembles the drug-resistant, *S. cerevisae* SF3B1 protein (**Fig. 1A**).

**Figure 1.**
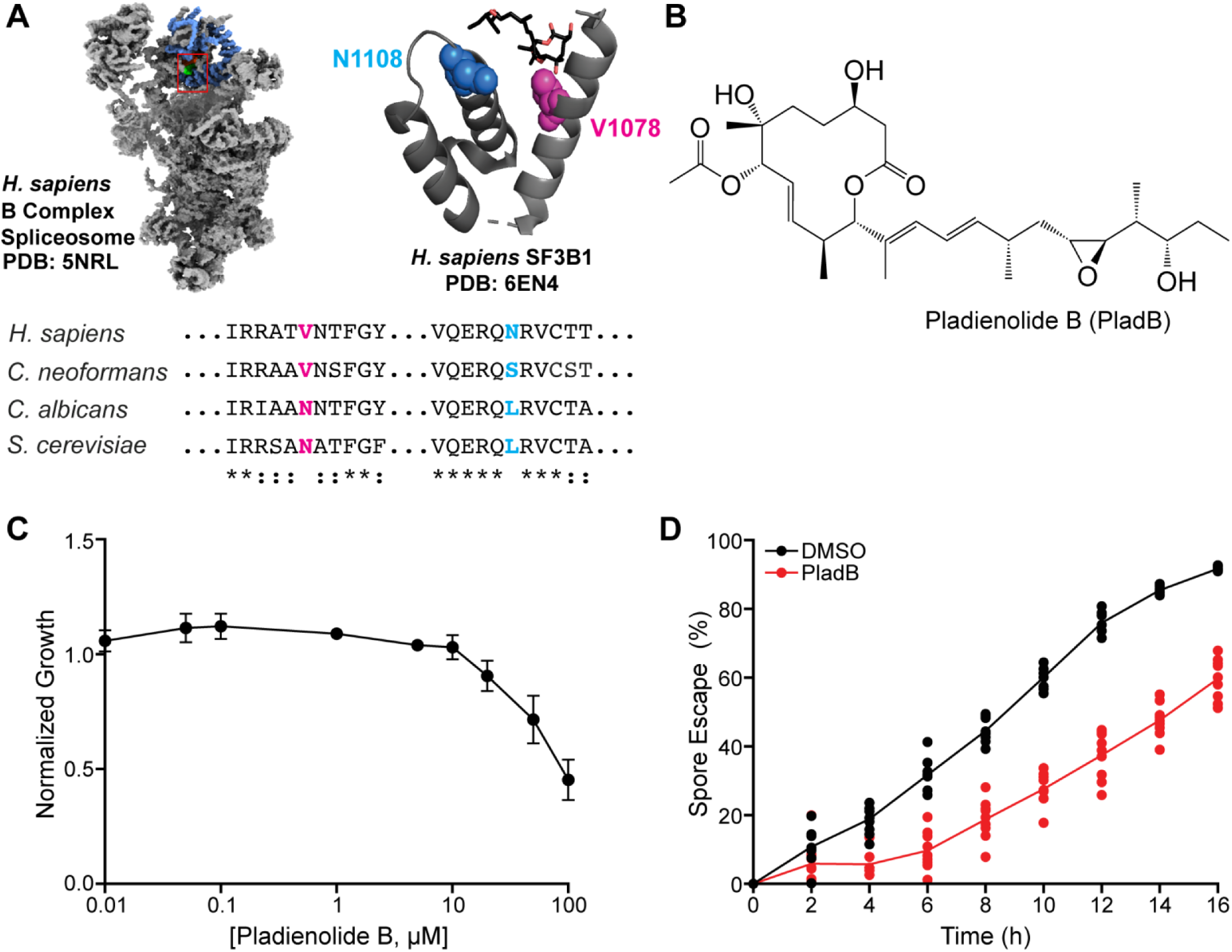
Pladienolide B Inhibits *C. neoformans* Yeast Growth and Spore Germination. (**A**) (Left) Structures of the human B complex spliceosome with SF3B1 highlighted in blue. The region of SF3B1 corresponding to the view in the image to the right is shown in a red box. (Right) The drug- and branch point-binding pocket of SF3B1 with key amino acids for resistance in *S. cerevisiae* shown in spacefill and pladienolide B (PladB) shown in stick representation. Protein sequences for *C. albicans* (strain SC5314) and *C. neoformans* (strain H99) were obtained from UniProt. **(B)** Chemical structure of pladienolide B (PladB). **(C)** Normalized growth of *C. neoformans* strain KN99α in liquid culture relative to a DMSO control in the presence of increasing concentrations of PladB (±SD from *N* = 3 biological replicates). **(D)** Percent spore escape during germination of *C. neoformans* serotype D spores (JEC20×JEC21) in the presence of DMSO or 30 µM PladB. Lines were fit to the mean values per time point from *N* = 9 technical replicates. Structures shown in panel A were generated using ChimeraX and Pymol (Mura et al., 2010; Pettersen et al., 2021).

To test for sensitivity of *C. neoformans* to SF3B1-binding splicing modulators, we treated *C. neoformans* strain KN99α with varying concentrations of pladienolide B (PladB, **Fig. 1B**) in liquid YPD medium and monitored yeast growth. We observed that growth slowed at PladB concentrations >10µM with 50% growth inhibition at 100 µM (**Fig. 1C**). In comparison with a drug-sensitized *S. cerevisiae* strain, PladB is ∼100-fold less potent in *C. neoformans* under these conditions for reducing fungal growth (Hansen et al., 2019). To determine if growth slowed due to cell death or failure of cells to reproduce, we collected yeast after 24 h of PladB treatment and observed robust growth on plates once the drug was removed (**Fig. S1**). This suggests that PladB, under these conditions, is fungistatic rather than fungicidal.

Germination is a critical step in the *C. neoformans* life cycle in which spores transition into vegetatively growing yeast. The differentiation of spores into yeast is required to cause disease in a host (Ortiz & Hull, 2024). Given the inhibitory effects of PladB on yeast growth, we next assessed whether the drug similarly impacts spore germination. Using a previously described quantitative germination assay (QGA) (Barkal et al., 2016; Frerichs et al., 2021), we monitored spore germination over 16 h in the presence or absence of PladB (30 µM). The QGA tracks changes in cell morphology to quantify the frequency and rate of transition from spores to yeast during germination. Consistent with the growth data, PladB reduced spore escape by ∼30%, indicating impaired germination (**Fig. 1D**). Thus, the splicing modulator PladB can affect *C. neoformans* by inhibiting both yeast growth and spore germination.

### PladB Potency Increases in Combination with FK506 or Clorgyline

In *S. cerevisiae*, sensitivity to SF3B1-binding splicing modulators can be observed only if drug efflux pumps are deleted, suggesting that efflux pumps contributed to resistance to the splicing modulators by decreasing their intracellular concentrations (Hansen et al., 2019). We hypothesized that a similar mechanism of resistance could account for the lower potency of PladB in *C. neoformans* relative to the sensitized *S. cerevisiae* strain. To test this hypothesis, we identified two compounds reported to inhibit ATP-binding cassette (ABC) and/or major facilitator superfamily (MFS) drug efflux pumps, which are often implicated in drug resistance (Chen et al., 2011; Chen et al., 2012; Fu et al., 2017; Holmes et al., 2016; Holmes et al., 2012; Kozubowski et al., 2008; Lima et al., 2019). Specifically, FK506 (**Fig. 2A**, top) has been reported to inhibit several fungal pleiotropic drug resistance (PDR-type) transporters (Tanabe et al., 2019). Similarly, clorgyline (**Fig. 2A**, bottom) synergizes with azole antifungals by inhibiting the Cdr1 and Mdr1 drug efflux pumps in *C. albicans* and *C. glabrata* (Toepfer et al., 2023).

**Figure 2.**
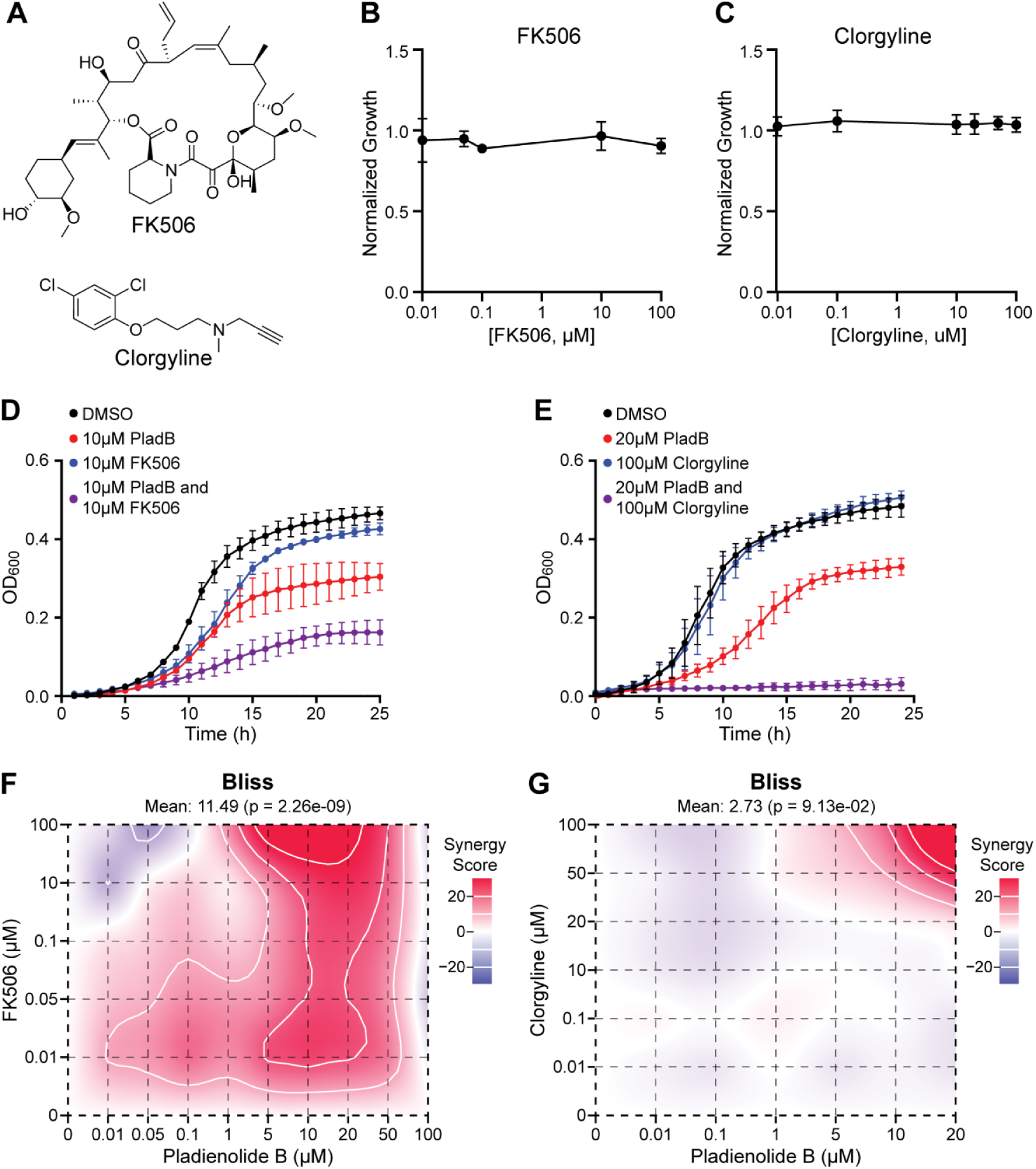
PladB Potency Increases in Combination with FK506 or Clorgyline. **(A)** Chemical structures of FK506 and clorgyline. **(B, C)** Normalized growth of *C. neoformans* strain KN99α in the presence of increasing concentrations (0.01–100 µM) of FK506 (B) or clorgyline (C) (±SD from *N* = 3 biological replicates). **(D, E)** Growth curves of *C. neoformans* strain KN99α in the presence of DMSO, PladB alone, FK506 alone, Clorgyline alone, or combinations of PladB with FK506 (D) or clorgyline (E) (error bars represent the ±SD from *N* = 3 replicates). **(F, G)** 2D contour maps depicting the combinatorial effects of PladB with FK506 (F) and clorgyline (G) in *C. neoformans* strain KN99α, where red and blue represent areas of high or low synergy values, respectively, calculated using the Bliss synergy scoring method. In panels B-E, data points were connected by straight lines for clarity.

Based on these reports, we tested whether sensitivity to PladB in *C. neoformans* could be increased by addition of either FK506 or clorgyline. We first evaluated the effects of these drugs in the absence of splicing modulation by measuring *C. neoformans* growth and discovered that neither FK506 nor clorgyline significantly impacted growth at concentrations up to 100 µM (**Fig. 2B, C**). Despite minimal impact of FK506 or clorgyline alone on yeast growth, *C. neoformans* exposed to combinations of these drugs with PladB grew very poorly (**Fig. 2D, E**). Addition of FK506 (10 µM) reduced yeast growth an additional 2-fold in the presence of PladB (10 µM), while addition of clorgyline (100 µM) together with PladB (20 µM) led to full growth inhibition.

To further evaluate the impact of these drug combinations, we co-varied the concentrations of PladB and FK506 or clorgyline and calculated the extent of growth inhibition by measuring the optical density of each culture after 24 h (**Fig. S2**). We then analyzed the data using SynergyFinder Plus (Zheng et al., 2022) software and the Bliss independence model for drug action, which assumes that drugs act independently through distinct mechanisms to produce their combined effects (Bliss, 1939; Yadav et al., 2015). We chose this model for its ability to evaluate drugs with non-overlapping targets, as is likely the case for PladB and FK506 or clorgyline.

For PladB, this analysis revealed a large degree of synergy at PladB concentrations > 5 µM and nearly all concentrations of FK506 (**Fig. 2F**). This was reflected in the mean synergy score for the assay of 11.49, reflecting synergistic effects across a range of concentrations. In the presence of FK506, the potency of PladB greatly increased and approached that of the drug-sensitized *S. cerevisiae* strain (∼50% growth inhibition at 1-10 µM PladB). In contrast, clorgyline had relatively low synergy scores across most conditions (mean score of 2.73) (**Fig. 2G**), indicating that at these concentrations PladB and clorgyline largely had additive effects. Exceptions occurred at the very highest concentrations of clorgyline (50-100 µM), which displayed synergies with high concentrations of PladB (20 µM; **Fig. 2G**, upper right corner). Together, these results show that FK506 and clorgyline can increase the potency of the splicing modulator PladB for inhibiting *C. neoformans* growth in liquid culture and that these effects can be synergistic.

### PladB in Combination with FK506 or Clorgyline Blocks Spore Germination

To determine whether or not FK506 or clorgyline could increase the inhibition activity of PladB on the ability of spores to germinate, we used QGAs to evaluate germination efficiency.

Under control conditions (DMSO alone), most spores germinated and transitioned to yeast after 16h (**Fig. 3A**, **B**; data column 1). In the presence of FK506 (50 µM) or clorgyline (50 µM) alone, germination also proceeded normally, and similar numbers of spores, intermediates, and yeast were observed relative to the DMSO control after 16 hours of germination. As described above (**Fig. 1D**), treatment with PladB (30 µM) alone partially inhibited the transition from spores to yeast (**Fig. 3A**, **B**; data column 4). In striking contrast, however, combining PladB with either FK506 or clorgyline completely blocked the transition from spores to yeast (**Fig. 3A**, **3B**; data columns 5 and 6). Similar to the yeast growth results, FK506 and clorgyline can increase the potency of PladB as a spore germination inhibitor.

**Figure 3.**
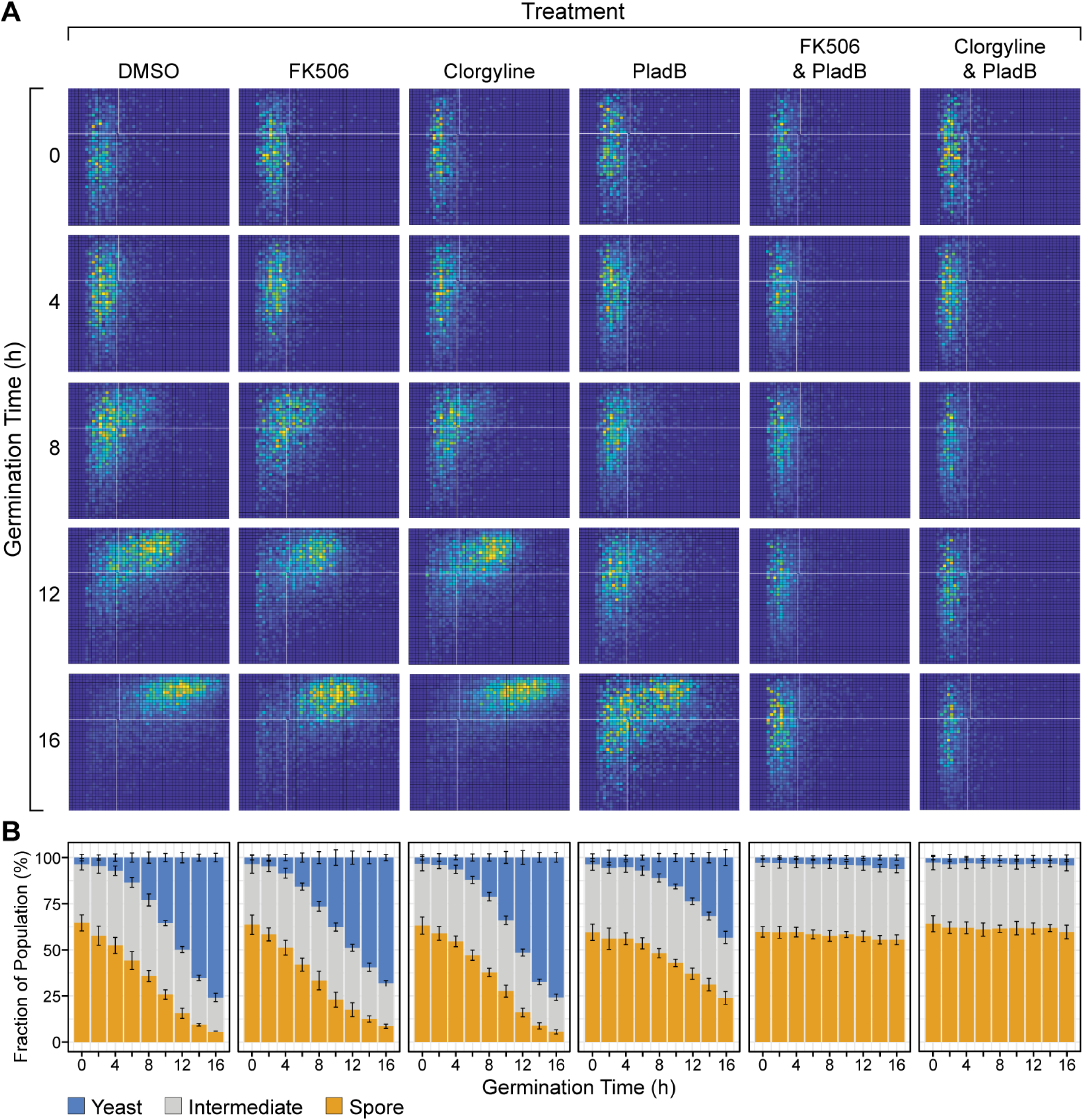
PladB in Combination with FK506 or Clorgyline Inhibits *C. neoformans* Spore Germination. **(A)** Germination profiles of *C. neoformans* serotype D spores (JEC20×JEC21) in the presence of DMSO, FK506 (50 µM), clorgyline (50 µM), PladB (30 µM), or their combinations. Each two-dimensional histogram was obtained by quantitative analysis of microscopy data where each x-axis represents cell area (µm^2^) and each y-axis represents cellular aspect ratio (width/length). Pixel intensities represent the number of cells with particular values of area and aspect ratio. Spores have smaller areas and aspect ratios (lower left) relative to yeast (upper right). **(B)** Stacked bar plots quantifying cell populations at each time point from the histograms shown in panel (A). (Error bars represent ±SD from *N* = 9 technical replicates). Portions of the data shown for the PladB condition here were also used to generate the plot in Fig. 1D.

### FK506, PladB, and Their Combination Have Unique Impacts on Gene Expression

We next determined whether growth inhibition of *C. neoformans* in the presence of PladB was due to pre-mRNA splicing inhibition and if changes in the transcriptome could provide insight into the synergistic effects of PladB and FK506. We focused on FK506 rather than clorgyline in order to assay PladB at lower concentrations (10 µM) when growth inhibition is minimal in the absence of the other drug (FK506; also 10 µM). Under these conditions, PladB and FK506 have a high calculated synergy value (23.1).

We collected total RNA from yeast (KN99α) after exposure to 1% (v/v) DMSO (solvent control), 10 µM PladB, 10 µM FK506, or both PladB and FK506 at 10 µM for 2 h. Paired-end RNA-Seq (40-50 million reads/sample) was then used to identify the isolated RNAs after polyA selection. The resultant reads were filtered and aligned to the *C. neoformans* genome. Principle component analysis indicated that the three replicates for each condition clustered together and were segregated from the other samples. This indicates similarities in the transcriptomes between replicates and that each condition resulted in a distinct transcriptomic signature (**Fig. 4A**).

**Figure 4.**
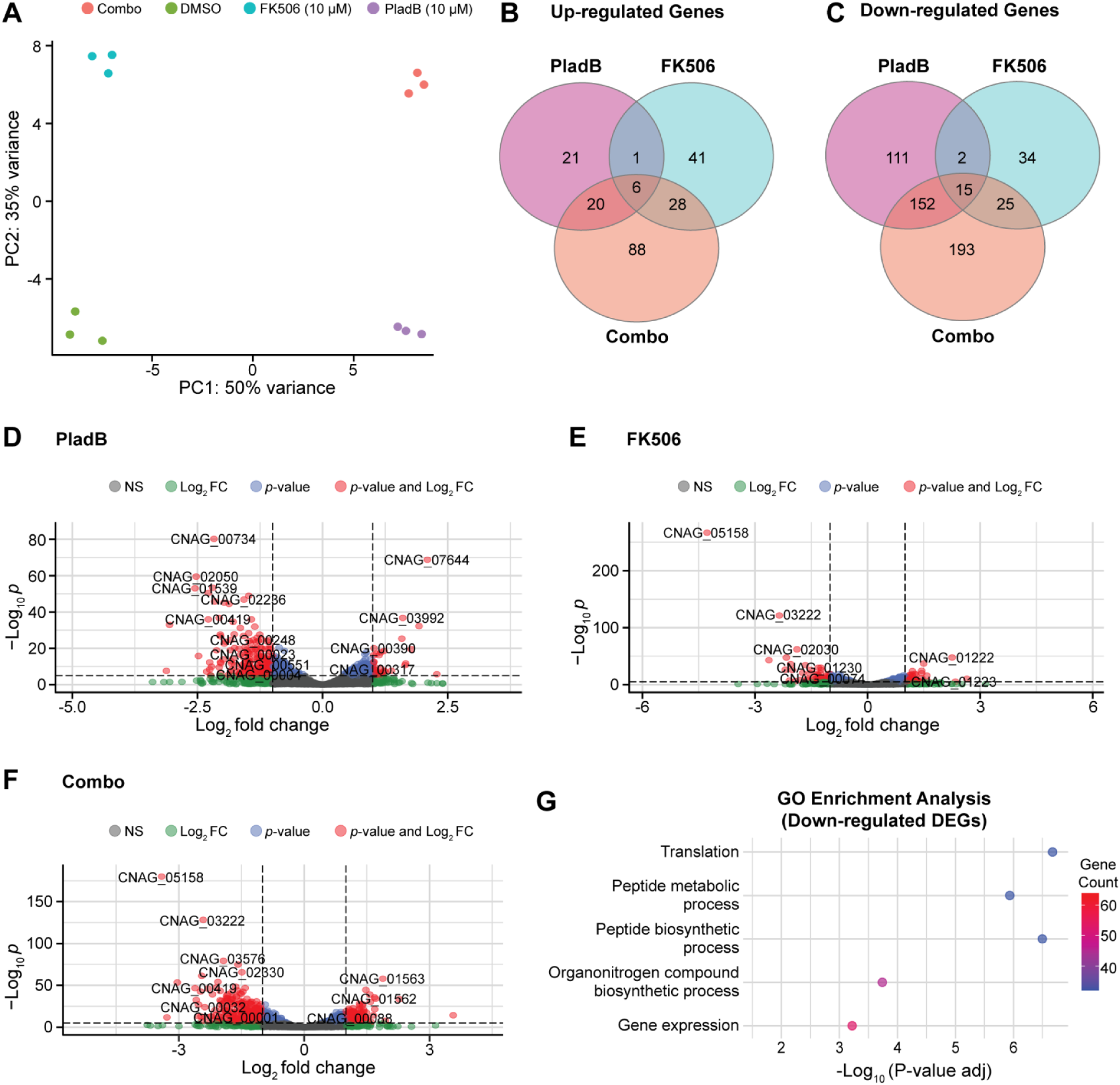
FK506, PladB, and Their Combination Have Unique Impacts on Gene Expression. **(A)** Principal component analysis (PCA) of RNA-seq data illustrating variance among conditions and clustering of replicates (*N* = 3) for cells treated with either DMSO or 10 µM of either PladB or FK506 or their combination (Combo). **(B**, **C)** Venn diagrams illustrating common and unique up- and down-regulated (panels B and C, respectively) differentially expressed genes (DEGs) in each condition conditions. **(D–F)** Volcano plots displaying DEGs in *C. neoformans* strain KN99 α treated with PladB (panel D), FK506 (panel E), or the combination of PladB and FK506 (panel F). DEGs were defined as those with log₂ fold change <-1 or >1 and a -log₁₀ *p*-value >0.05 (red dots). 6979 genes (not including any non-coding RNAs) were analyzed in each plot. **(G)** Gene ontology (GO) enrichment analysis of downregulated DEGs in the combination treatment of PladB and FK506.

We conducted a differential gene expression analysis to identify up-regulated and down-regulated differentially expressed genes (DEGs) in each condition relative to the DMSO control (**Fig. 4B**, **C**). While some genes were shared among all conditions, each drug treatment also resulted in unique sets of DEGs. FK506 treatment alone resulted in the fewest DEGs, consistent with its minimal impact on growth (**Fig. 2B**). In contrast, the combination treatment of FK506 with PladB resulted in the largest number of DEGs, also consistent with growth inhibition under these conditions.

Among the DEGs, it is interesting to note that one of the most highly down-regulated genes in yeast exposed to FK506 or FK506 with PladB is the *C. neoformans* intron lariat debranching enzyme (CNAG_03222) (**Fig. 4D-F**). This gene was also down-regulated in the presence of PladB alone, but to a lesser extent than when FK506 was present. Lariat debranching is a critical step in pre-mRNA processing because it allows for turnover of the excised intron and may facilitate the release of splicing factors (Buerer et al., 2024). It is possible that accumulation of intronic RNA due to decreased expression of debranchase contributes to the increased sensitivity of *C. neoformans* to PladB under synergistic conditions with FK506.

It is also worth noting that when yeast are exposed to both FK506 and PladB, genes involved in mitochondrial stress response show changes in expression, with cytochrome C assembly protein (CNAG_03576) downregulated and prohibitin PHB1 (CNAG_00088) upregulated **(Fig 4F)**. The downregulation of cytochrome C assembly suggests a disruption in mitochondrial respiration, potentially impairing ATP production and increasing oxidative stress (Guaragnella et al., 2012; Kim et al., 2013; Su et al., 2014). Meanwhile, the upregulation of PHB1, a mitochondrial chaperone, may indicate a compensatory stress response aimed at stabilizing mitochondrial function and preventing apoptosis (Nijtmans et al., 2000; Signorile et al., 2019; Steglich et al., 1999). This opposing regulation suggests that the combination of PladB and FK506 induces mitochondrial stress, which may also contribute to fungal growth inhibition.

A gene ontology (GO) analysis of all of the DEGs was carried out for each condition. This analysis failed to detect significant enrichment among any particular biological process for the up-regulated DEGS. However, the GO analysis revealed strong enrichment for translation factors among the down-regulated DEGs, specifically when both PladB and FK506 were present (**Fig. 4G**). It is likely that down regulation of the translational machinery also plays a major role in growth inhibition under these conditions.

### PladB Inhibits pre-mRNA Splicing in *C. neoformans*

Given the central role of pre-mRNA splicing in gene regulation and its impact on nearly all expressed genes in *C. neoformans* (Gonzalez-Hilarion et al., 2016; Grützmann et al., 2014; Janbon, 2018), we hypothesized that the drug treatments would induce changes in intron processing. Visual inspection of the data indicated that while few intronic reads were observed in samples treated with DMSO or FK506, intronic reads could be more readily observed in the presence of PladB (see **Fig. 5A**, as an example).

**Figure 5.**
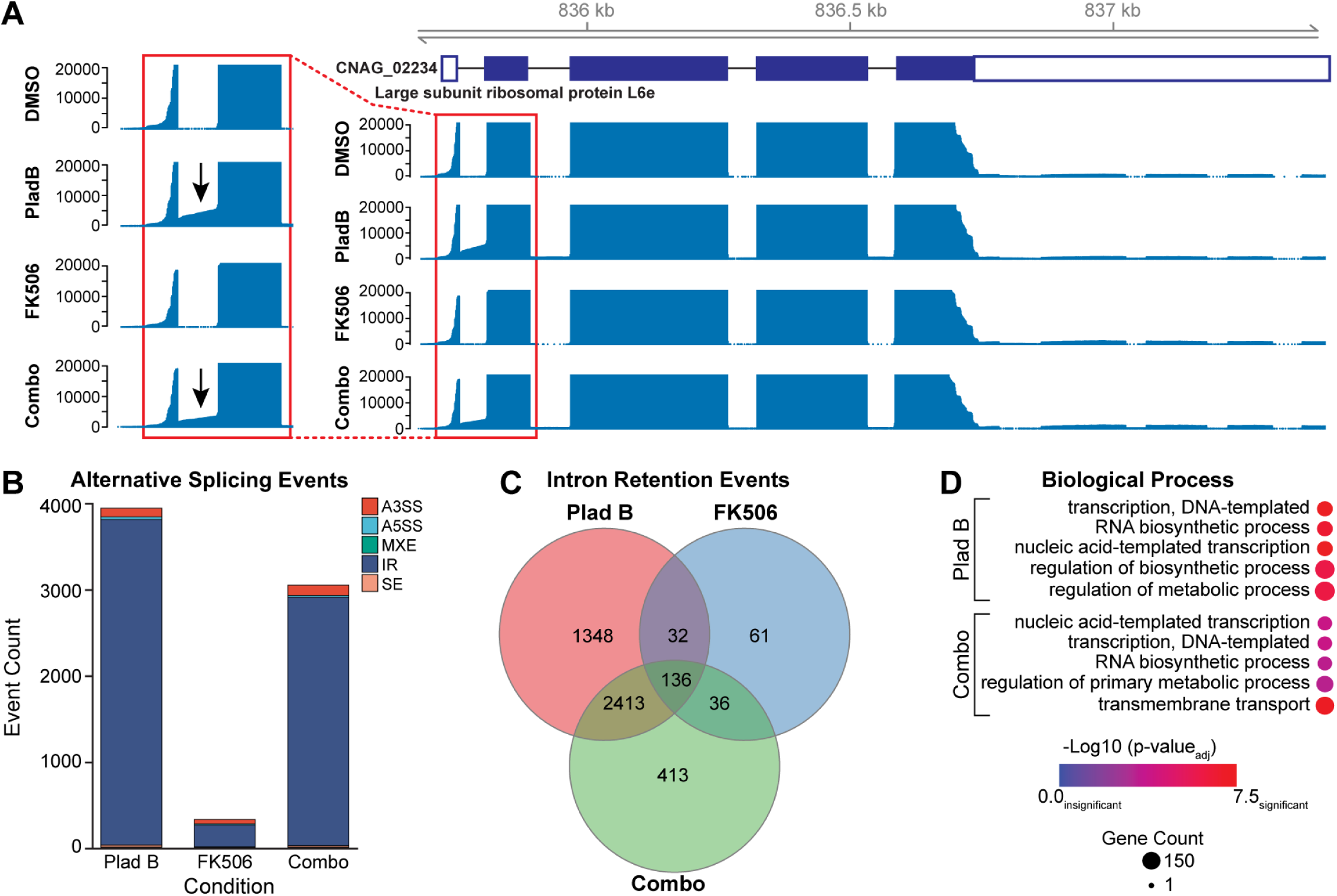
PladB Inhibits pre-mRNA Splicing in *C. neoformans.* **(A)** Example read coverage tracks over the gene CNAG_02234 (encoding the large subunit ribosomal protein L6e) in the presence of DMSO, PladB, FK506, or the combination of PladB with FK506. Note the presence of intronic reads in PladB-treated conditions (arrows). **(B)** Stacked bar graph depicting the distribution of alternative splicing events across each condition. A3SS and A5SS represent alternative 3’ and 5’ splice site selection, respectively; MXE represents mutually exclusive exon usage, IR represents intron retention, and SE represents skipped exons. **(C)** Venn diagram showing the numbers of unique and common intron retention events under each condition relative to the DMSO controls. **(D)** Gene ontology (GO) enrichment analysis of genes with retained introns in PladB-containing conditions.

To assess changes in splicing specifically, we used SpliceWiz to detect differential alternative splicing events based on a percent spliced-in (PSI) metric (Wong et al., 2023). We filtered the detected changes in splicing to those resulting in a change in PSI of more than 0.1 (10%) and a false discovery rate (FDR) of less than 0.05. Importantly, PSI was calculated from reads spanning intron/exon junctions, not just the intronic reads. Therefore, changes in expression of debranchase (**Fig. 4D-F**) and lariat intron accumulation are unlikely to account for changes in PSI.

Using these metrics, we identified thousands of changes in splicing in the presence of PladB, nearly all of which resulted in intron retention (**Fig. 5B**). This is consistent with PladB functioning as a splicing modulator in *C. neoformans*, and the most frequent outcome of this modulation being failure to remove the intron (retention). We detected more intron retention events when PladB was present alone than when in combination with FK506, suggesting that increased numbers of retained introns alone cannot account for the increased growth inhibition observed due the presence of FK506. It is also worth noting that we observed intron retention in only a subset of all introns (∼10% of the >40,000 introns found in *C. neoformans*) (Janbon, 2018). This is consistent with work carried out both in *S. cerevisiae* and in human cells, showing that different introns have differential sensitivities to splicing modulators (Corrionero et al., 2011; Hunter et al., 2024). It is possible that the full extent of intron accumulation cannot be discerned from these experiments because RNA surveillance pathways such as nonsense mediated decay (NMD) may degrade many of the transcripts with retained introns.

Comparison of the intron retention events between samples showed that (as with the DEG analysis) each condition resulted in its own unique set of retained introns with some introns being retained across multiple conditions (**Fig. 5C**). We then considered the possibility that introns retained only in the PladB with FK506 condition could share some unique features that led to retention only in the presence of both drugs. *C. neoformans* intron architecture is noteworthy because most introns tend to be small (average of 65bp) relative to those in humans or *S. cerevisiae* (Janbon, 2018). When analyzed by intron size, the introns uniquely retained in the presence of PladB or PladB with FK506 had average lengths closely matching the genome-wide intron average (**Fig. S3A**). Intron retention also did not seem to be related to intron order (*e.g.,* specifically impacting the first or last introns in transcripts) as the distributions of intron retention events vs. intron ordinal number seemed quite similar for each condition (**Fig. S3B**). We do note, however, that these steady-state results could be biased due to mRNA decay pathways (or other processes) and the accumulation of partial transcripts (for example, 3′◊5′ decay by the exosome could contribute to the larger number of observed intronic reads corresponding to first, 5′-most introns). Analysis of the 5′ and 3′ splice sites of the retained introns showed no significant deviation from the consensus sequences of all introns (**Fig. S3C**), suggesting that these sequences also do not play an obvious role in their retention.

Finally, GO analysis of the genes with retained introns showed no enrichment for any particular biological process for RNAs isolated from yeast grown in the presence of FK506, which showed few intron retention events overall. However, PladB-containing conditions showed that many of the retained introns were detected in transcripts originating from genes involved in transcription, RNA biosynthesis, and metabolic process regulation (**Fig. 5D**). These results suggest that pre-mRNA splicing modulation by PladB in *C. neoformans* can slow growth by particularly impacting a subset of cellular processes involving RNA production itself.

## DISCUSSION

In this work, we have shown that PladB can inhibit growth of *C. neoformans* in liquid culture as well as spore germination. These inhibitory activities likely stem from retention of introns in many pre-mRNAs, confirming that PladB functions as a splicing modulator in *C. neoformans*. These results also confirm that splicing activity is essential for germination. Both FK506 and clorgyline increase the potency of PladB inhibition. At certain concentrations, these drugs act synergistically with PladB to potently block yeast growth and spore germination. Transcriptomic analysis shows that together, FK506 and PladB impact gene expression and pre-mRNA splicing.

The mechanism of synergy between PladB and FK506 (and at high concentrations, clorgyline) could arise from inhibition of drug efflux pumps and increased intracellular concentrations of PladB. However, it is interesting to note that we did not observe significantly more intron retention events when PladB and FK506 were used in combination relative to PladB alone. If higher intracellular concentrations of PladB do result from FK506, it is possible that synergy arises from modulation of a different repertoire of splicing events (**Fig. 5C**) rather than overall increased levels of intron retention. An important limitation of this hypothesis is that we did not measure changes in drug efflux due to FK506 or clorgyline directly, and we do not know the precise molecular target(s) of these compounds. Alternatively, FK506 causes significant down-regulation of a key enzyme involved in pre-mRNA splicing: the lariat intron debranchase. An alternate mechanism of synergy could arise from a block in lariat intron degradation that augments the action of the splicing modulator.

Growth inhibition due to PladB likely stems from preventing removal of a subset (∼10%) of *C. neoformans* introns. We have not yet been able to define what features of the impacted introns make them susceptible to PladB or not. This could involve both *cis* (intronic sequence) and *trans*-acting elements (transcriptional speed, chromatin environment) unique to each intron. DEGs and retained introns observed when PladB is used in combination with FK506 are enriched in processes including transcription, RNA biosynthesis, and translation. These results suggest that growth inhibition, and possibly spore germination inhibition as well, stems at least in part from disruption of core steps in gene expression.

Splicing modulators, including PladB and related compounds, have the potential to be potent antifungals able to arrest *C. neoformans* cell growth as well as prevent spore germination and initial infection. One limitation of PladB as an antifungal is that it also strongly inhibits the human splicing machinery (Effenberger et al., 2017). It will be interesting to see if fungal pathogen-specific splicing modulators, that do not inhibit human SF3B1 can be synthesized in the future. Recent work on derivatives of another splicing inhibitor (meayamycin) that also interacts with SF3B1 at the same site as PladB provides evidence that it may be possible to develop allele- or ortholog-specific drugs (Beard et al., 2024). Such fungal-specific splicing inhibitors based on PladB or meayamycin may also be co-administered with drug efflux inhibitors to increase their potency, as we have shown here with FK506 and clorgyline. Interestingly, FK506 is already used as an immunosuppressant (tacrolimus) for organ or tissue transplantation, and many of these same patients are at the greatest risk for fungal infection. It could be advantageous to develop splicing modulator antifungals that work synergistically with FK506 or other drugs that patients are already prescribed.

## AUTHOR CONTRIBUTIONS

SLL, MCM, CMH, and AAH: project conceptualization; JCP: media and reagent preparation; SLL, HV: yeast growth assays and analysis; MCM, NO, OV, SCO: spore germination assays and analysis; SLL: RNA isolation and RNA-seq analysis; MCM: RNA-seq genome alignment; SLL, MCM, CMH, and AAH: writing and editing of the manuscript.

## ACKNOWLEDGEMENTS

All authors were funded by a Badger Challenge program grant from UW-Madison. SLL, HV, and AAH were additionally funded by a EvansMDS Discovery research grant from the Edward P. Evans Foundation. SLL and MCM were funded by the Genetics Training Program (NIH 5T32GM007133). This work was also supported by an NIH R01 (AI137409) to CMH. We thank the UW-Madison small molecule screening facility for instrument access.

